# MetaCompass: Reference-guided Assembly of Metagenomes

**DOI:** 10.1101/212506

**Authors:** Victoria Cepeda, Bo Liu, Mathieu Almeida, Christopher M. Hill, Sergey Koren, Todd J. Treangen, Mihai Pop

## Abstract

Metagenomic studies have primarily relied on *de novo* approaches for reconstructing genes and genomes from microbial mixtures. While database driven approaches have been employed in certain analyses, they have not been used in the assembly of metagenomes. Here we describe the first effective approach for reference-guided metagenomic assembly of low-abundance bacterial genomes that can complement and improve upon *de novo* metagenomic assembly methods. When combined with *de novo* assembly approaches, we show that MetaCompass can generate more complete assemblies than can be obtained by *de novo* assembly alone, and improve on assemblies from the Human Microbiome Project (over 2,000 samples).

## Background

Microorganisms play an important role in virtually all of the Earth’s ecosystems, and are critical for the health of humans [1], plants, and animals. Most microbes, however, cannot be easily grown in a laboratory [2]. The analysis of organismal DNA sequences obtained directly from an environmental sample (a field termed metagenomics), enables the study of microorganisms that are not easily cultured. Metagenomic studies have exploded in recent years due to the increased availability of inexpensive high-throughput sequencing technologies. For example, the MetaHIT consortium generated about 500 billion raw sequences from 124 human gut samples in its initial analysis [3], and the Human Microbiome Project (HMP) has generated hundreds of reference microbial genomes and thousands of whole metagenome sequence datasets from healthy subjects [4].

The analysis of these vast amounts of data is complicated by the fact that reconstructing large genomic segments from metagenomic reads is a formidable computational challenge. Even for single organisms, the assembly of genome sequences from short reads is a complex task, primarily due to ambiguities in the reconstruction that are caused by genomic repeats [5]. In addition, metagenomic assemblers must be tolerant of non-uniform representation of genomes in a sample as well as of the genomic variants between the sequences of closely related organisms. Despite advances in metagenomic assembly algorithms over the past years [6–10], the computational difficulty of the assembly process remains high and the quality of the resulting assemblies requires improvement.

Consequently, many analyses of metagenomic data are performed directly on unassembled reads [11–15], however the much shorter genomic context leads to lower accuracy [16].The need for effective and efficient metagenomic assembly approaches remains high, particularly since long read technologies (which partly mitigate the challenges posed by repeats [17–19]) are not yet effective in metagenomic applications due to lower throughput, higher costs [20, 21], and higher required DNA quality and concentration.

Reference-guided, comparative assembly approaches have previously been used to assist the assembly of short reads when a closely related reference genome was available [22, 23]. Comparative assembly works as follows: short sequencing reads are aligned to a reference genome of a closely related species, then their reconstruction into contigs is inferred from their relative locations in the reference genome [23]. This process overcomes, in part, the challenge posed by repeats as the entire read (not just the segment that overlaps within adjacent reads) provides information about its location in the genome.

Currently, thousands of bacterial genomes have been sequenced and finished [24], and this number is expected to grow rapidly soon thanks to long read technologies. These sequenced genomes provide a great resource for performing comparative assembly of metagenomic sequences. Comparative approaches developed in the context of single genomes cannot, however, be directly used in a metagenomic setting. Simply mapping a set of reads to even hundreds of different genomes is currently computationally prohibitive. Furthermore, genome databases comprise many variants of a same genome (e.g., the US FDAs GenomeTrackr project [25] alone has contributed over 60,000 different strains of *Salmonella*), and genome by genome analyses would result in redundant reconstructions of metagenomic sequences. We also note that some recent reference-guided strategies implemented in genomic analysis tools, such as the “--trusted-contigs” feature of the SPAdes assembler [26, 27] and StrainPhlan [28] ignore the fact that the data being reconstructed originates from genomes that are related but different from the genomes found in public databases. As a result, such approaches may actually mis-assemble the metagenomic data exactly within the genomic regions where novel biological signals may be located.

In this paper, we describe the first effective assembly software package for the reference-assisted assembly of metagenomic data. We rely on an indexing strategy to quickly construct sample-specific reference collections, thereby dramatically reducing the computational costs of mapping metagenomic reads to references. Furthermore, we eliminate redundancy in the assembly by disambiguating the mapping of reads against closely related genomes, and identify differences between the metagenomic data and the reference genomes in order to reduce the likelihood of misassembly.

We show that our approach effectively complements *de novo* assembly methods. We also show that the combination of comparative and *de novo* assembly approaches can boost the contiguity and completeness of metagenomic assembly, and provide an improved assembly of the entire whole-metagenome sequencing data generated by the Human Microbiome Project [4].

Our software is released freely under an open-source license at:

http://www.github.com/marbl/MetaCompass.

## Results

All assemblies were compared based on contiguity statistics, number of errors, and based on the number of complete phylogenetic marker genes found in the final assembly – a measure of how useful an assembly may be to downstream analyses. The coverage of the set of marker genes has been used by the HMP and others [3, 29, 30] as measure of the completeness of an assembly.

### Evaluation of performance on synthetic metagenomic dataset

We first evaluated MetaCompass by assembling a synthetic microbial community [31]. The synthetic sample was downloaded from the NCBI Short Read Archive (SRA) database, (SRR606249) and contains 54 bacteria and 10 archaea. Among these organisms, 55 had complete genome sequences in the NCBI RefSeq database (the database used by default by MetaCompass), and 9 were available only as a high-quality draft assembly at the time of publication. Since the true genome sequences are known, these data are ideal as they allow us to fully quantify the quality of the genomic reconstruction.

We set the minimum coverage in MetaCompass at 1-and 2-fold (see Methods), then performed reference genome selection (see Methods and Supplementary Table 1). The assembly results (Table 1, see MetaCompass 1X and 2X) can be considered an approximate upper bound on the performance of any assembly tool, as in this case 90% of the genomes recruited were exactly those from which the metagenomic reads were obtained. We compared the performance of MetaCompass with that of three widely used *de novo* assemblers: IDBA-UD (July 2016) [8], MEGAHIT (v1.0.6) [32], and metaSPAdes (v3.9.0) [33]. Compared with these assemblers, MetaCompass achieved higher genome recovery (Table 1, Figure 1) and produced significantly larger and more accurate contigs (Table 1). When we decreased the MetaCompass minimum coverage threshold from 2-fold to 1-fold, we observed gains in maximum contig size and total aligned length, while retaining a similar error profile. However, we observe higher genome recovery at minimum coverage threshold 2 and 3. On the basis of the maximum contig size, total aligned length, error profile and genome recovery, we chose 2X as default setting for MetaCompass. Note that here we are not trying to prove that MetaCompass is better than *de novo* assemblers, and in this setting, the comparison is not fair because our reference collection contains the exact genomes present in the samples. Rather, we are trying to show that the performance of MetaCompass can be excellent if the reference collection contains genomes highly similar to those in the metagenomic sample being assembled.

**Figure 1.**
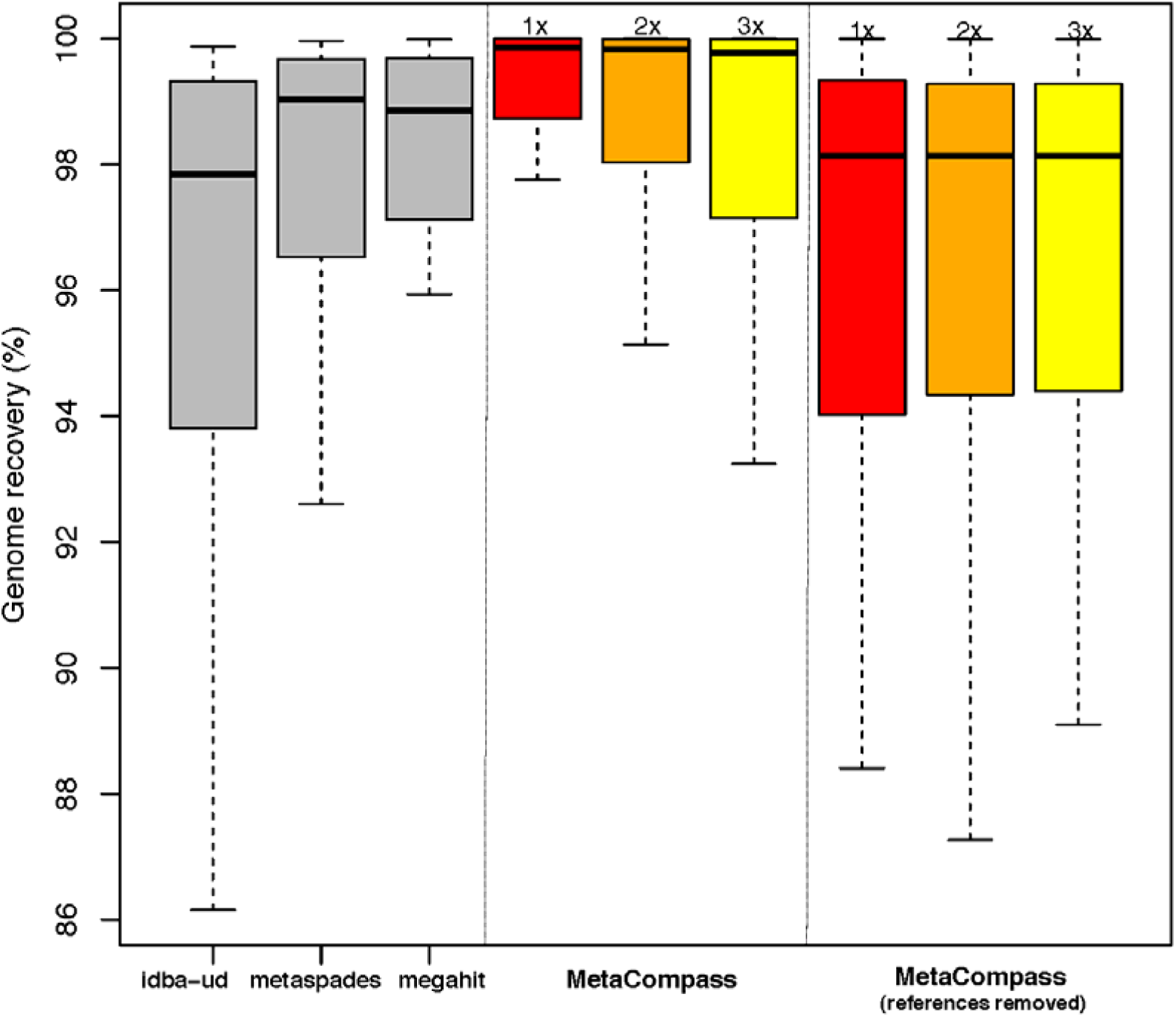
Genome recovery percentages in synthetic metagenome. (MetaCompass versus *de novo* assembly). Box plots represent distribution of genome recovery percentages (for the 64 genomes present in the synthetic metagenome). x-axis indicates the assembly method, either IDBA-UD, metaSPAdes, MEGAHIT, or MetaCompass. MetaCompass was run both with the reference genomes present in the database (recruited as described in the methods) and without the truth reference genomes in the database (they were individually removed). y-axis indicates the genome recovery percentage, 0% indicates the genome was unassembled, whereas 100% indicates the genome was fully assembled.

**Table 1.**
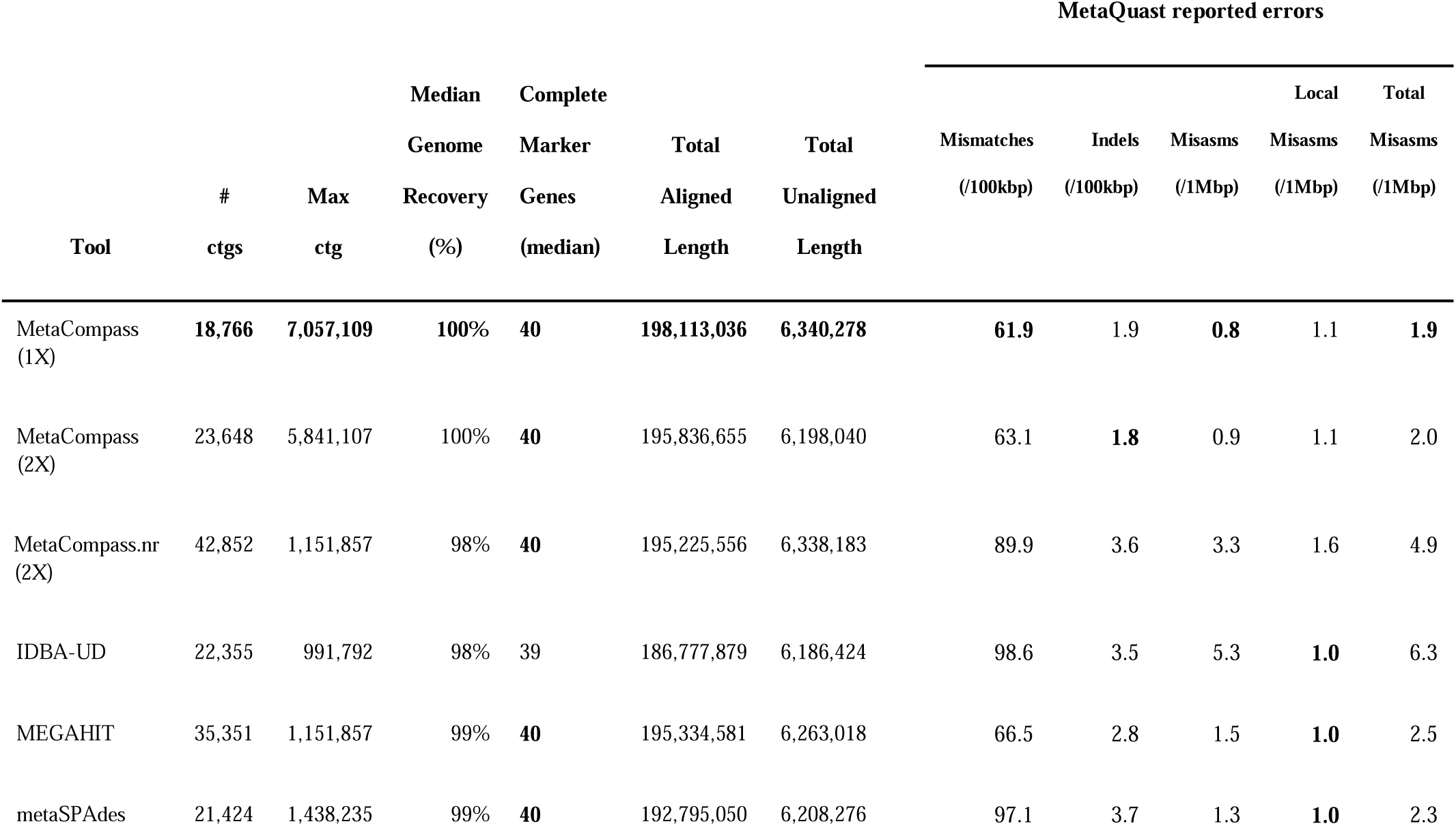
Evaluation of performance on synthetic dataset. MetaCompass (X) indicates the minimum coverage setting (1X or 2X), and MetaCompass.nr indicates all 64 reference genomes comprising the Shakya et al. dataset were removed from the database. **Tool** indicates the assembler, **# ctgs** the total number of assembled contigs reported by each assembler, **Max ctg** is the maximum contig length (broken at errors) for all assembled contigs, **Median Genome Recovery** (**%**) is the median percentage of each of the synthetic genomes that is recovered, **Complete Marker Genes (median)** is the median number of fully reconstructed marker genes, **Total Aligned Length** is the sum of the length of contigs aligned to the reference genomes, **Total Unaligned Length** is the sum of the length of unaligned contigs, and **MetaQuast reported errors** are error statistics generated with MetaQuast.

### References removed from database

To provide a better idea of how MetaCompass would perform in a worst-case scenario, we removed from the database the genomes represented in the mock community (Supplementary Table 2), thereby forcing MetaCompass to recruit near-neighbor reference genomes, when available. (see ‘MetaCompass.nr’ row, Table 1). Median genome recovery for MetaCompass is just 1% less than that of *de novo* assemblers. The accuracy of the reconstruction, as measured by mismatch and indel rates, is higher than that of IDBA-UD and metaSPAdes (Table 1, MetaCompass.nr (2x)), while moderately lower than MEGAHIT.

The number of misassemblies and local misassemblies per 1 Mbp of assembled sequence (as reported by MetaQuast [34]) increased from 2.0 to 4.9 when reducing the coverage threshold to 1. To put this increase into context, we measured the total number of possible errors by evaluating the "accuracy" of the near-neighbor reference genomes recruited by MetaCompass with respect to the correct reference sequence (Figure 2, see hashed blue bar). This allows us to capture the real differences between the recruited reference genomes and the actual genome represented in the synthetic dataset [31], essentially providing an upper bound for the number of errors MetaCompass would make if it simply recapitulated the sequence of the selected reference genomes. As seen in Figure 2, MetaCompass is making five times fewer errors than would be expected, indicating our software is not unduly biased by the sequence and structure of the reference genome.

**Figure 2.**
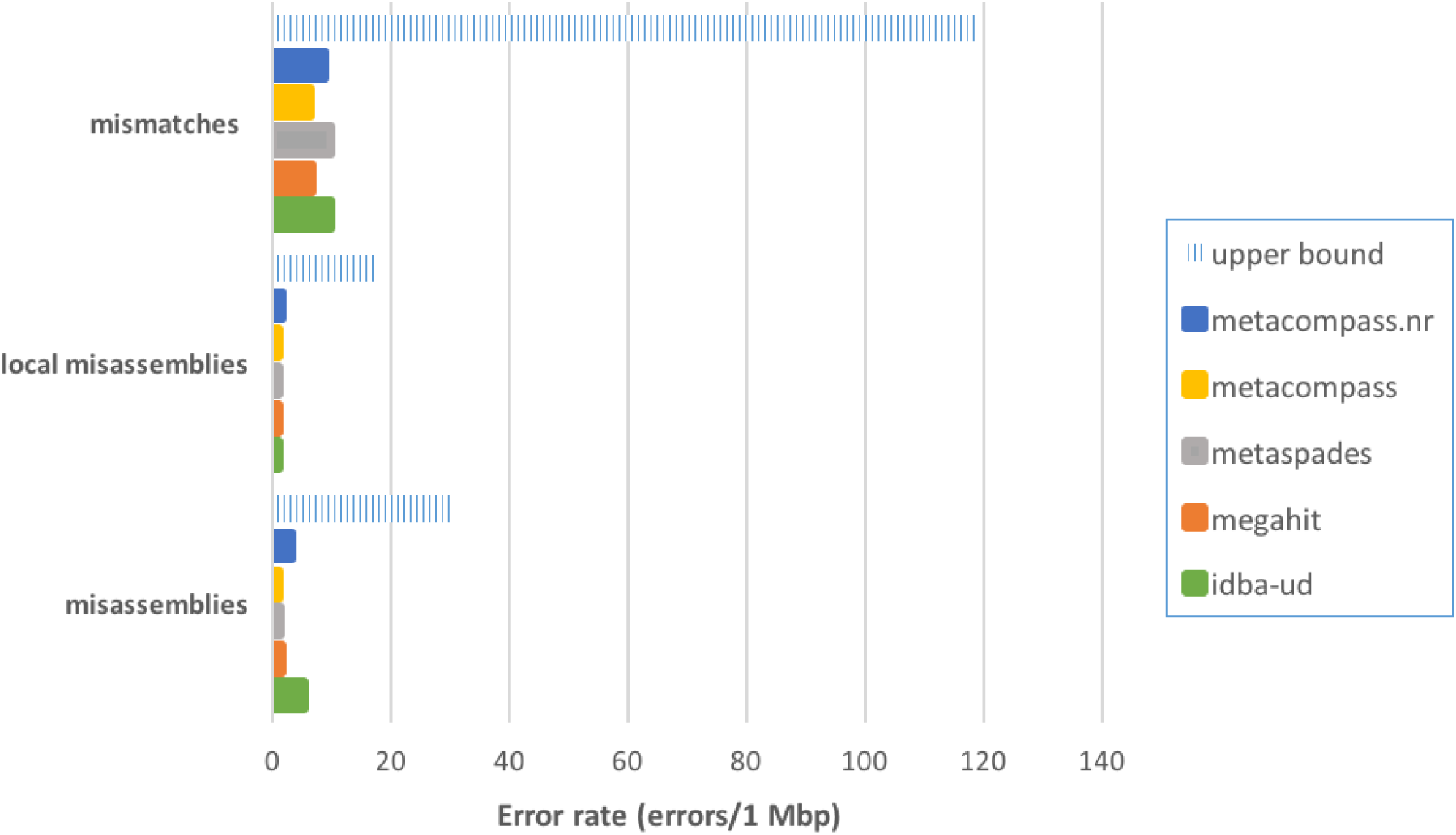
Error profile on synthetic dataset. The hashed blue bar represents the difference between the second-best reference genome (recruited by MetaCompass) and the true genome represented in the sample. This bar can be viewed as an upper bound on the errors metacompass.nr could make if it simply reconstructed the reference genome. **Mismatches** are the number of bases in a contig that differ from the reference genome. **Misassemblies** include large-scale (left flanking region aligns >1 kbp away from right flanking region) relocations, interspecies relocations, translocations, and inversions. **Local misassemblies** include small-scale (left flanking region aligns <=1 kbp away from right flanking region) translocations and inversions. All errors are normalized to represent rates per 1 Mbp.

### Evaluation of performance on downsampled synthetic datasets

To evaluate the ability of MetaCompass to assemble low-coverage genomes, we down-sampled the synthetic dataset to just 5 million paired-end reads, or 10% of the original data set. After downsampling, the average coverage was reduced to approximately 3-fold (data not shown). The results (Table 2, Figure 3) highlight that MetaCompass can recover a median of 90% of each of the 64 genomes in the sample. While metaSPAdes comes in second place and is able to recover 80% (median recovery), it does so at the cost of a four times higher mis-assembly rate. The two remaining methods, MEGAHIT and IDBA-UD, leave a quarter to a half of the genomes unassembled (Table 3).

**Figure 3.**
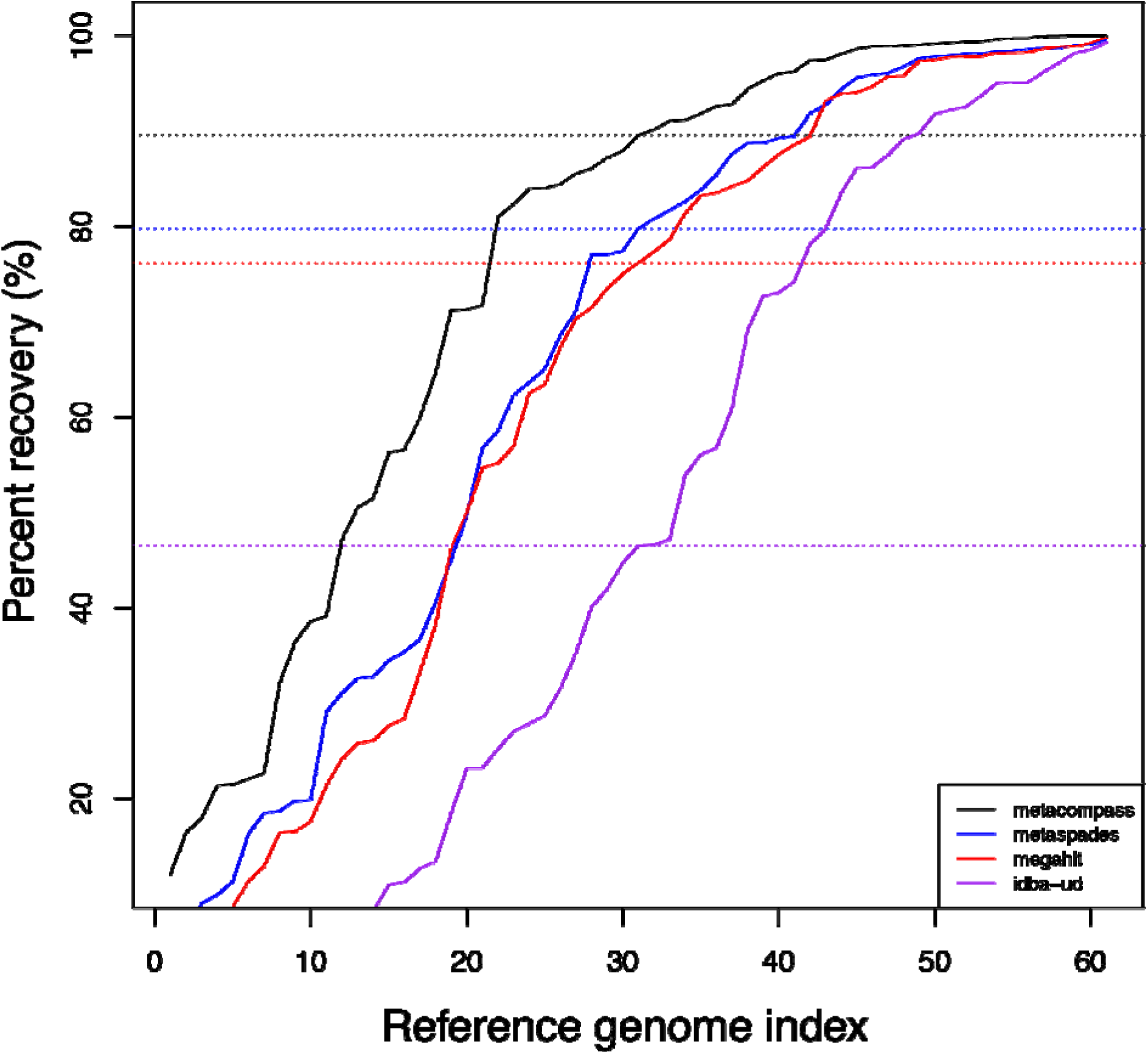
MetaCompass performance on low coverage dataset. Results obtained by downsampling the Shakya et al. synthetic genome to just 10% of the original set of reads. The 64 genomes present in the sample are ordered per assembler by percent recovery, from lowest to highest. The y-axis indicates how much of the n-th reference was covered by correctly assembled contigs (can range from 0% to 100%). The colored dashed lines indicate the median percent recovery for each assembler.

**Table 2.** Evaluation of performance on down-sampled synthetic dataset. The synthetic dataset was down-sampled to only contain 10% of the total reads. **Tool** indicates the assembler, **# ctgs** the total number of assembled contigs reported by each assembler, **Max ctg** is the maximum contig length (broken at errors) of all assembled contigs, **Median Genome Recovery** (**%**) is the median percentage of each of the synthetic genomes that is recovered, **Complete Marker Genes (median)** is the median number of fully reconstructed marker genes, **Total Aligned Length** is the sum of the length of contigs aligned to the truth genomes, **Total Unaligned Length** is the sum of the length of unaligned contigs, and **MetaQuast reported errors** are error statistics generated with MetaQuast [54].

|  |  |  |  |  |  |  | MetaQuast reported errors |  |  |  |
| --- | --- | --- | --- | --- | --- | --- | --- | --- | --- | --- |
| Method | #<br>ctgs | Max<br>Ctg | Median | Complete | Total<br>Aligned<br>Length | Total<br>Unaligned<br>Length | Mismatches | Indels | Misasms | Misasms |
|  |  |  | Genome<br>Recovery<br>(%) | Marker<br>Genes<br>(median) |  |  | (/100kbp) | (/100kbp) | (>1 kbp) | (<1 kbp) |
| MetaCompass | 71457 | 962,929 | 90% | 22 | 134,008,055 | 3,009,931 | 117.6 | 1.9 | 112 | 33 |
| IDBA-UD | 43973 | 120159 | 45% | 6 | 75,970,693 | 1,564,008 | 175.0 | 5.3 | 3447 | 93 |
| MEGAHIT | 62842 | 209,706 | 76% | 15 | 105,665,678 | 2,774,432 | 128.0 | 4.1 | 772 | 122 |
| metaSPAdes | 67138 | 287,554 | 80% | 16 | 111,636,826 | 3,154,199 | 133.0 | 4.3 | 470 | 115 |

**Table 3.**
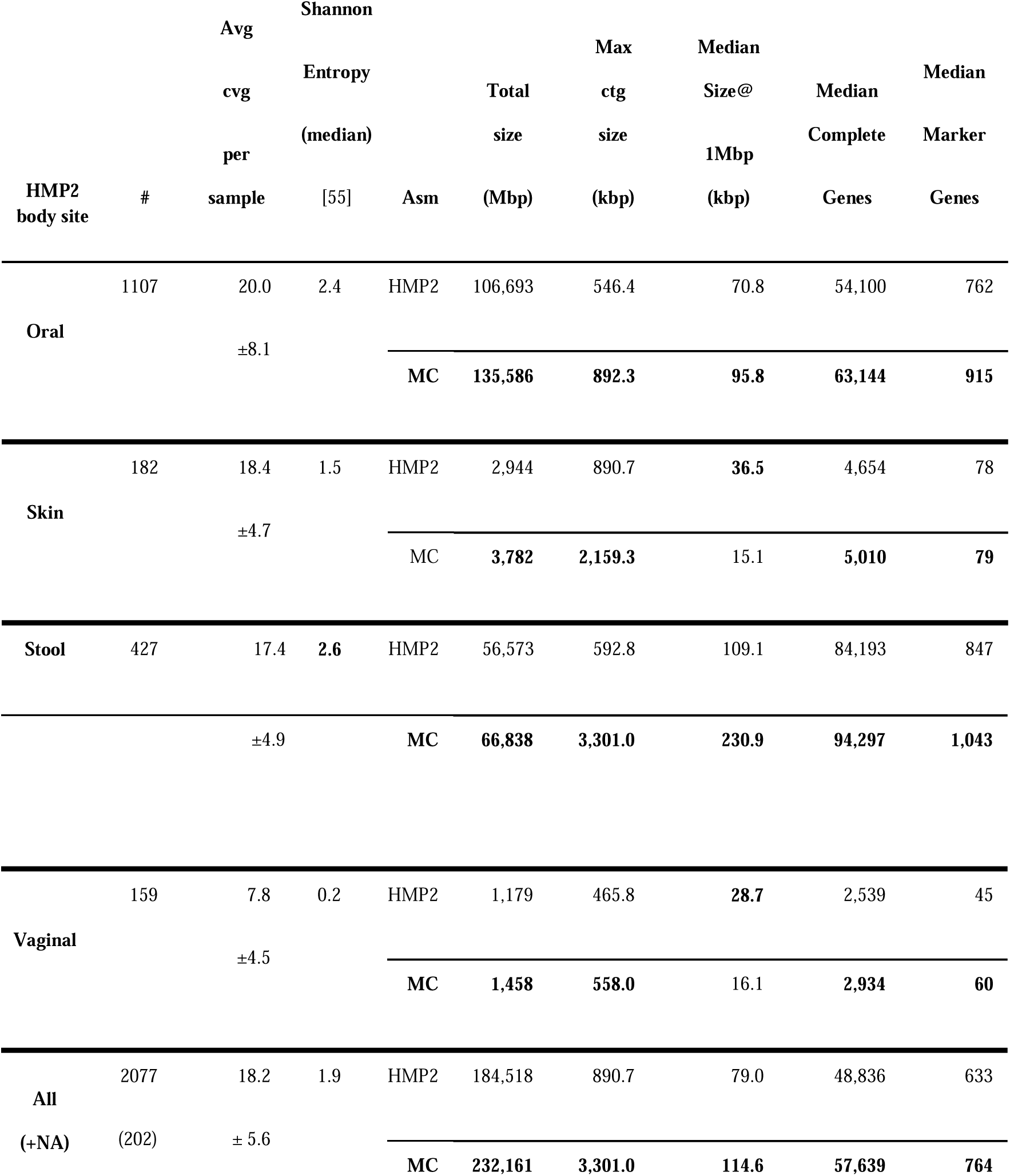
Re-assembly of 2,077 samples generated in the Human Microbiome Project. The results are aggregated by body site. **#** indicates the total reads per sample, **Avg cvg per sample (X)** is the mean estimate read coverage calculated based on the de novo assembly of each sample and body site, **Shannon Entropy (median)** is the Shannon diversity value per body site as reported in Li *et al* 2012 [55].The rows labeled MC contain results obtained with MetaCompass. The rows labeled HMP2 show the statistics for contigs from the production HMP2 assembly. **Total Size** (Mbp) is the total assembly size for each method, **Max ctg size (kbp)** is the size of the largest contig, **Median Size@1Mbp (kbp)** represents the mediad size of the largest contig C such that the sum of all contigs larger than C exceeds 1Mbp. **Median Complete Genes** represents the median number of complete genes per sample. **Median Marker Genes** indicates the median number of complete marker genes per sample.

### Computational performance

When dealing with large-scale data sets, the combination of total required memory and run time is an important factor in determining the applicability of a computational tool. We first evaluated the runtime performance of MetaCompass on a Linux 12-core server node with 80 GB of memory using the Shakya et al. synthetic dataset. The wall clock run time on this synthetic dataset for MetaCompass is comparable to that of *de novo* assemblers, sometimes lower (Supplementary Table 3). MetaCompass (without PILON) and Megahit were the only approaches that required <16GB of RAM on a 100 million read dataset, highlighting the scalability of these methods to large datasets.

### Reassembly of the data generated by the Human Microbiome Project (HMP2)

To further explore the benefits and limits of comparative approaches for metagenomic assembly, we re-analyzed with MetaCompass 2,077 metagenomic samples from the HMP Project (ftp://public-ftp.hmpdacc.org/Illumina/PHASEII/). These samples cover 15 body sites from four broad regions of the human body: oral, skin, stool, and vaginal. We compared the assemblies produced by MetaCompass with the official IDBA-UD assemblies reported by the HMP project [35]. Note that these assemblies were recently improved by Lloyd-Price et al. [36] but we did not include them in this study. Across all samples, on average, MetaCompass outperforms the HMP2 *de novo* approach, leading to an overall better assembly of the original data (Table 3, Figure 4). However, the relative performance of MetaCompass and the HMP2 assembly varied across body-sites due to the specific characteristics of the microbial communities being reconstructed. While MetaCompass generates more assembled sequence and complete marker genes across all body sites, the maximum contig size and size at 1 Mbp metrics vary per body site. In oral and stool samples (Figure 4), MetaCompass outperforms *de novo* assembly for all metrics. In skin and vaginal samples (Figure 4), the *de novo* approach has better contiguity statistics but MetaCompass assembles more complete marker genes. To gain further insight into these results we calculated the average nucleotide identity between the *de novo* assembled contigs and the recruited reference genomes for each body site. In all body sites, except for oral, the assembled contigs had 99% average nucleotide identity to the reference genomes. In the oral samples, the most distant reference genomes had only 97% identity to the assembled contigs, indicating that at least in part, the lower effectiveness of MetaCompass is due to the absence of a sufficiently closely related reference genome for some of the oral samples.

**Figure 4.**
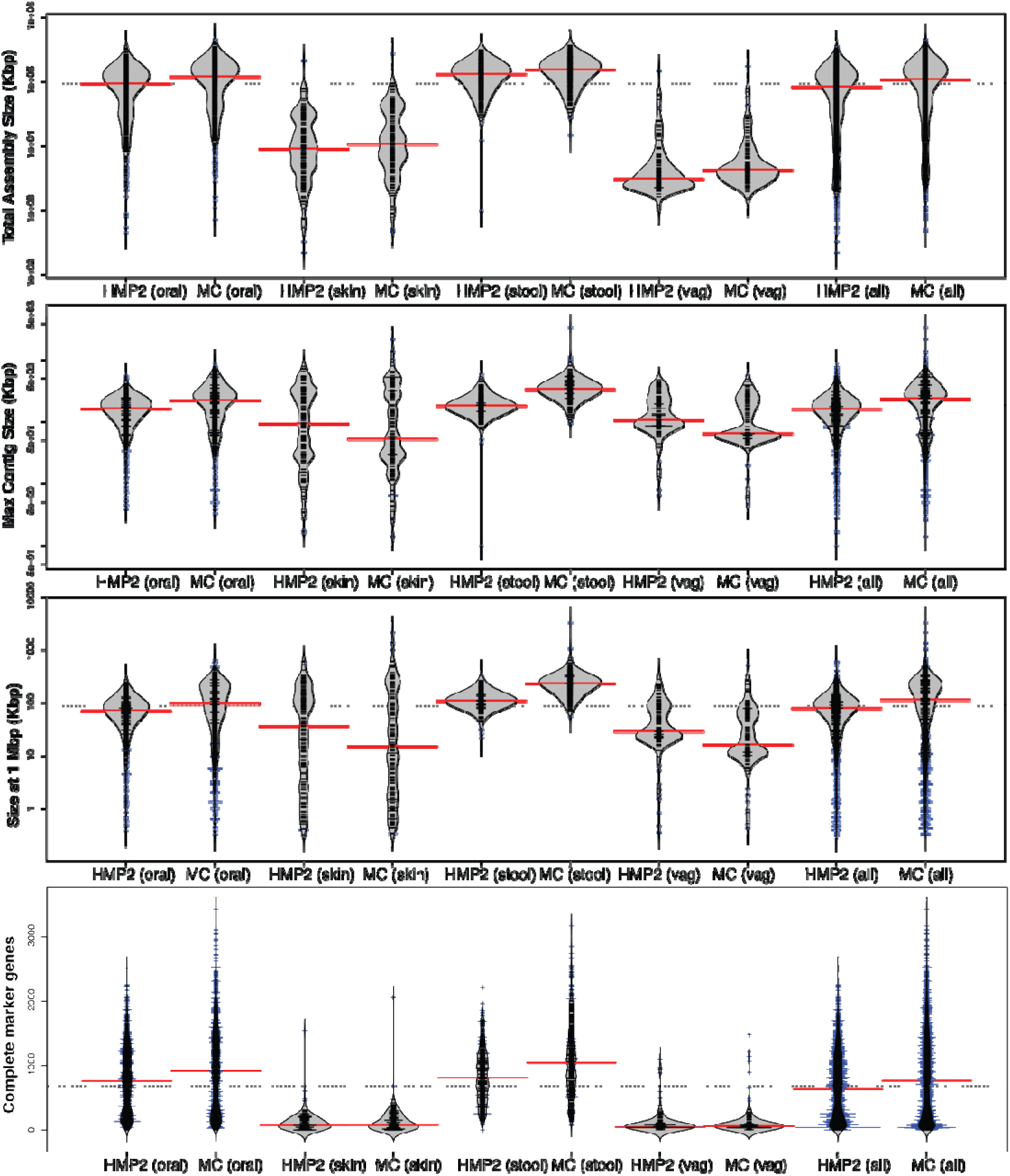
Comparative assembly of 2,077 metagenomic samples from the HMP2 Project. The ‘bean plots’ represent the distribution of assembly contiguity and completeness statistics across all samples within the data. The x axis organizes the data by assembly and body site. The y-axis indicates the statistic used to evaluate the assembly contiguity or completeness. The top panel shows total assembly size (kbp), the second panel shows maximum contig size (kbp), the third panel shows the size of the contig at 1 Mbp, and the bottom panel shows the complete marker genes assembled per sample.

To further explore the drop in contiguity in skin and vaginal samples, we focused on just the contigs that mapped to bacterial genomes contained in the reference database, allowing for a direct comparison between MetaCompass and *de novo* contigs. The results shown in Table 4 indicate that for this set of contigs, MetaCompass outperforms the *de novo* approach for the vaginal samples. However, the *de novo* HMP2 assembly of the skin sample is still better in terms of complete genes recovered, but equivalent to MetaCompass with respect to complete marker genes recovered (a measure of assembly completeness).

**Table 4.**
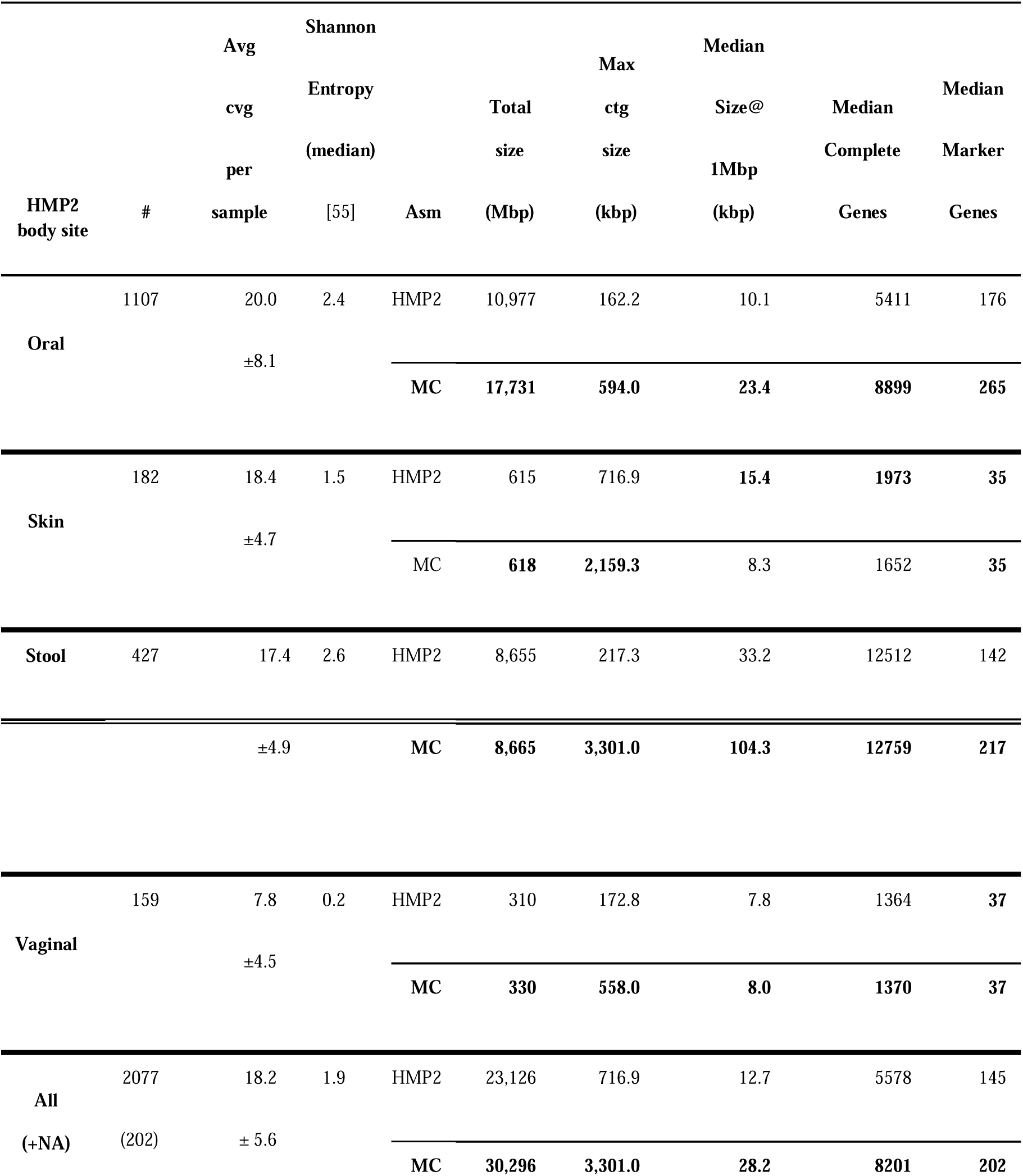
Results of Human Microbiome Project analysis within the reference genomes. The results are aggregated by body site. **#** indicates the total reads per sample, **Avg cvg per sample (X)** is the mean estimate read coverage calculated based on the de novo assembly of each sample and body site, **Shannon Entropy (median)** is the Shannon diversity value per body site as reported in Li *et al* 2012 [55].The rows labeled MC contain results obtained with MetaCompass. The rows labeled HMP2 show the statistics for contigs from the production HMP2 assembly. **Total Size** (Mbp) is the total assembly size for each method, **Max ctg size (kbp)** is the size of the largest contig, **Median Size@1Mbp (kbp)** represents the median size of the largest contig C such that the sum of all contigs larger than C exceeds 1Mbp. **Median Complete Genes** represents the median number of complete genes per sample. **Median Marker Genes** indicates the median number of complete marker genes per sample.

### Comparing reference-guided to *de novo* assembly on low-coverage HMP2 samples

To assess the ability of MetaCompass to assemble low-abundance organisms, we focused on all skin HMP2 samples. The skin samples had the second lowest average number of reads while still containing reasonable diversity and richness, as reported in Table 3. We removed the contigs assembled via *de novo* assembly from the MetaCompass output, collected the reference genomes that were used, mapped the HMP2 contigs to these reference genomes, and then evaluated the number of complete genes and complete marker genes in both. Compared to the HMP2 assembly, referenceguided assembly of these low coverage samples is able to reconstruct approximately 10% more marker genes (4,423 versus 3,915) than the *de novo* approach, roughly equating to 10 additional complete bacterial genomes in total.

We next searched for microbes that were present in the skin samples at relatively low coverage and explored the differences between the reconstructions generated by the HMP2 project and MetaCompass. Specifically, we identified a low coverage assembly of a *Propionibacterium acnes* genome reconstructed by both MetaCompass and the HMP in sample SRS057083. The HMP2 assembly covers less than 40% of the closest reference genome (NC_016516.1, *Propionibacterium acnes* TypeIA2 P.acn33), while the MetaCompass assembly covers more than 90% of the same genome.

## Discussion

The benefit of comparative assembly is highly dependent on the reference genomes available in the database provided to MetaCompass. While MetaCompass can effectively use reference genomes that are distantly related to the genomes being assembled, the quality of the reconstruction is lower than can be achieved with closely related reference sequences. As the set of genome sequences available in public databases continues to increase, so will the effectiveness of reference-guided assembly approaches such as MetaCompass.

We have shown MetaCompass to be particularly effective in the assembly of low coverage or rare microbes, setting in which *de novo* assembly approaches simply cannot be used with good results. Improved assembly of low-abundance, rare microbes from existing datasets has the potential to provide valuable information in complex microbial communities or clinical samples where the host DNA comprises a large fraction of the data. Clinical applications are also a particularly relevant application domain for comparative approaches as the vast majority of publicly available genome sequences comprises human pathogens.

While MetaCompass provided an advantage over *de novo* approaches for most of the humanassociated microbial communities sampled by the HMP project, in skin samples the performance of MetaCompass was on average lower than the assemblies produced by the HMP. This result could be due to structural genome dynamics of bacterial defense systems commonly found in skin microbes [37–39], situation that introduces frequent structural variants between the reference genomes and the corresponding environmental isolates. We plan to further explore this hypothesis through graphbased analyses of *de novo* assemblies of the corresponding communities.

MetaCompass relies on the taxonomic profiling tool MetaPhyler as an efficient indexing strategy for identifying the reference genomes most closely related to the data being assembled. Compared to whole-genome indices, the MetaPhyler index is based on just 18 phylogenetic marker genes that are ubiquitous in bacteria, thus providing a compact and efficient data-structure. Using marker genes ensures that any genome present at a high enough coverage to allow assembly will be detected despite indexing just a small fraction of its genome. Since MetaPhyler, and other similar tools [40, 41] are designed for much broader use cases than those targeted here, it is likely that better performance in both memory and speed can be achieved by an indexing strategy designed specifically for comparative metagenomic assembly, and we plan to explore such strategies in future work. Furthermore, comparative assembly provides new opportunities for the development of sequence alignment approaches that optimize the combined time of index creation and alignment. Most of the recent developments in sequence alignment have assumed index construction to be a one-time off-line operation, trading off a computationally intensive indexing approach for more efficient queries.

## Conclusion

We have described MetaCompass, a computational pipeline for comparative metagenomic assembly. This novel method for metagenomic assembly leverages the increasing number of genome sequences available in public databases. We have shown that comparative and *de novo* assemblies provide complementary strengths, and that combining both approaches effectively improves the overall assembly, providing a consistent increase in the quality of the assembly. Even when distant reference genomes are recruited, MetaCompass is competitive with *de novo* genome assembly methods. These results are due to two critical steps. First, reference bias is avoided by constructing the consensus sequence from the reads within the sample, using the reference genome as just a guide, and by breaking the assembly where the reads indicate a structural disagreement with the reference. Second, unmapped reads are used in a *de novo* assembly process to reconstruct the sections of the metagenomic sample that are not similar to known reference genomes. In summary, we believe that reference-guided approaches such as MetaCompass, will increasingly replace the more computationally expensive and error-prone *de novo* assembly approaches as the collection of available reference genome sequences increases.

## Methods

### Methods overview

First, we use MetaPhyler [13] to identify reference genomes that are most closely related to the data represented in the input a sample. We use the NCBI RefSeq genome database (June 2016) as the standard reference collection for MetaCompass. We only retain for further consideration the genomes estimated by MetaPhyler to be represented at sufficient depth of coverage. These genomes are aligned using Bowtie2 [42] (v2.2.9). The resulting read alignments are then used to identify a minimal set of genomes that best explain all read alignments, then the read alignments are used to construct contigs. We developed the tool buildcontig to generate a consensus sequence for the contigs and then use Pilon [43] (v1.18) to correct the contigs in a way that reflects the genome being assembled and to avoid biasing the reconstruction towards the reference sequence. Contigs may be broken at this stage if the metagenomic sequence diverges from the reference sequence. Finally the reads that were not included in the reference-guided process outlined above are assembled using MEGAHIT [32] (v1.0.6) to reconstruct the metagenomic segments not represented in the reference collection. The details of each analysis step are described below.

### Selecting reference genomes

While comparative assembly approaches have already been described for single genomes [23, 44] their use in metagenomic data is complicated by the multiple unknown organisms and the thousands of genomes available in public databases. Building efficient indexes for large reference collections is computationally challenging for short read aligners [41], both in term of speed and memory consumption. For assembly, however we only need to use the genomes that are detected in a sample. To speed up the genome reference selection step, we reduce the 31 marker genes in MetaPhyler to 18 universally conserved marker genes in bacteria and archaea (intersection between the sets of genes used by FetchMG [45, 46] and MetaPhyler [13]). The MetaPhyler index is much smaller than a whole-genome index, yet still allows us to identify the closest reference genome detected in the sample being assembled. We further speed up the execution of MetaPhyler by restricting the analysis to just those reads that share at least one 28-mer with one of the marker gene sequences in the database. We rely on kmer-mask (http://kmer.sourceforge.net/wiki/index.php?Main_Page) to execute this filtering step. The selected reads are then aligned to the marker collection using BLASTN with the parameters ‘-word_size 28-evalue 1e-10-perc_identity 95-max_target_seqs 100’ and a minimum HSP alignment length of 35. Since closely related genomes can share the same marker genes, we retain all hits with a bit score equal to that of the top hit. Finally, we exclude from further consideration all the genomes with an estimated coverage below a user-selected coverage threshold (2-fold, by default).

### Aligning reads to reference sequences

The results presented in the paper are based on aligning the reads to the selected reference genomes with Bowtie 2 [42] (parameters: --sam-nohead --sam-nosq -- end-to-end --quiet --all-p 12). The alignments are then filtered to keep ties of lowest edit distance for each reads, allowing a read to be aligned in multiple locations similar to the best-strata option of bowtie1.

### Selecting a minimal reference set

In its simplest form, the comparative assembly approach involves mapping the reads to a genome and using their relative placement within this genome to guide the construction of contigs [23]. In the context of metagenomic data, however, this process is complicated by the fact that individual reads may map to multiple reference genomes, some of which are highly similar to each other. Adequately dealing with this ambiguity is critical for effective assembly. If all read mappings are retained, allowing a read to be associated with multiple reference genomes, the resulting assembly will be redundant, reconstructing multiple copies of the homologous genomic regions. If for each read a random placement is selected from among the multiple equivalent matches, none of the related genomes may recruit enough reads to allow assembly, thereby leading to a fragmented reconstruction. Assigning reads to genomes according to their estimated representation in the sample (determined, e.g., based on the number of reads uniquely mapped to each genome), may bias the reconstruction towards the more divergent reference genomes, which may lead to an overall poorer reconstruction of the genomic regions shared across related genomes. Here we propose a parsimony-driven approach – identifying the minimal set of reference genomes that explains all read alignments.

Formally, this problem can be framed as a set cover problem, an optimization problem which is NP-hard. To solve this problem, we use a greedy approximation algorithm, which iteratively picks the set of genomes that covers the greatest number of unused reads. It can be shown that this greedy algorithm is the best-possible polynomial time approximation algorithm for the set cover problem [47].

### Building contigs

Given a set of reference genomes, selected as described above, a set of shotgun reads, and the alignment between each read and reference genome, the process of creating contigs is straightforward. For each nucleotide in each reference genome, we look at the bases from the reads that are mapped to each locus, and pick the variant (nucleotide or indel) with the highest depth of coverage as the consensus and report it. Minimum depth of coverage and length for creating contigs can be specified through the program command-line options.

### Removing reference-bias with Pilon

Differences between the sequences being assembled and the reference genome used by MetaCompass can degrade the performance of the comparative assembly process. We employ Pilon [43] to "polish" the reference-guided assemblies, thereby changing the consensus sequence to resemble the data in the sample rather than the reference genome. During this process we also identify signatures of larger differences between the metagenomic sample and the reference sequence, and break the assembly at those locations.

### Combining reference-guided and de novo assembly

We employ the *de novo* assembler MEGAHIT to assemble reads that were unable to be mapped back to the reference-guided assembly generated by MetaCompass. These reads represent microbes that are missing from our reference database and novel variants. This approach allows the final assembly to capture both reference and non-reference sequences. We chose MEGAHIT because it is currently the most efficient *de novo* assembler for metagenomics [48]. MEGAHIT is also the default assembly methods for the JGI metagenomic pipeline [49] and performed well in a recent review [50].

### Gene prediction and marker gene detection

The genes were predicted in the contigs using MetaGeneMark [51](v3.26) with the “MetaGeneMark_v1.mod” model parameter file and using the option “-n” to remove partial genes containing long strings of “N”. The completion status of the genes (complete, lack 5’, lack 3’ and lack both) was defined by detecting all the common start codon (“ATG”, “TTG”, “GTG”) and stop codon (“TAA”, “TAG”, “TGA”) of prokaryotic genes. The 40 universal single copy marker proteins [52, 53] were identified in predicted genes using the standalone version of fetchMG (v1.0) http://www.bork.embl.de/software/mOTU/) [46].

### MetaQuast validation parameters

The command used to run MetaQuast was: ‘metaquast.py-R./shakya_references --fragmented --gene-finding’

### Synthetic metagenome assembly parameters

IDBA-UD requires a single fasta file that was generated using the IDBA ‘fq2fa --merge --filter’ command. MEGAHIT was run using the options ‘-- presets meta-sensitive --min-count 3 --min-contig-len 300-t 12’. MetaSPAdes was run using the options ‘--meta-t 12’, then all contigs shorter than 300nt and with less than 3X coverage were removed. IDBA-UD was run using the options ‘--min_count 3 --min_contig 300 --mink 20 --maxk 100 --num_threads 12’. MetaCompass was run using the options-m [1,2,3]-g 300-t 16’ on the synthetic dataset and ‘-m 3-g 300-t 16’ on the HMP2 samples.

### Data availability

A list of all available HMP samples was obtained by combining those available from the HMP Data Analysis and Coordination Center (DACC) (www.hmpdacc.org) and the HMP SRA project PRJNA48479 on 11/16/2016. Any sample listed in the SRA and not in the DACC was downloaded and processed by the HMP WGS Read Processing Protocol (http://www.hmpdacc.org/doc/ReadProcessing_SOP.pdf). Three DACC samples were corrupt or extracted to a duplicate SRS identifier (SRS023176, SRS043422, and SRS057182) and were downloaded from SRA and processed as above. A total of 98 454 samples were excluded from the downloaded set. This resulted in 2,713 samples. Some samples (504) had no references recruited and were excluded from further analysis. This resulted in 2,209 MetaCompass assemblies. All HMP2 assemblies available at ftp://public-ftp.hmpdacc.org/HMASM/IDBA/ <u>were downloaded (2,341 total assemblies).</u> A total of 2,077 samples (Supplementary Table 4) had both an HMP2 assembly and a MetaCompass assembly and were used for the analysis.

The set of known genomes for the synthetic dataset is available via the Supplementary Table 2 from Shakya *et al* [31].

### Software availability

MetaCompass is available as an open-source package at: https://github.com/marbl/MetaCompass. The code is licensed under the Artistic License 2.0: https://opensource.org/licenses/Artistic-2.0

## List of abbreviations

**DACC –** Data Analysis and Coordination Center

**HMP –** Human Microbiome Project

**NCBI –** National Center for Biotechnology

**RefSeq -** NCBI Reference Sequence Database

**SRA** – Short read archive

## Funding

The authors were supported in part by the NIH, grants R01-HG-004885 and R01-AI-100947, by the NSF, grants IIS-1117247 and IIS-0812111, and the Office of Naval Research under cooperative agreement number N00173162C001, all to MP. SK was supported by the Intramural Research Program of the National Human Genome Research Institute, National Institutes of Health. SK has received funding for travel and accommodation expenses in order to speak at Oxford Nanopore Technologies conferences.

## Author’s contributions

BL and MP designed the approach. VC, BL, TT, and MA implemented the algorithms, methods, and scripts described in the paper. VC, TT, and SK generated the assemblies presented in the paper. TT, VC, and MA validated the assembly results and performed the comparisons between different assemblers. VC, MP, BL, TT and wrote the paper. All authors were involved in reviewing and revising the manuscript. All authors read and approved the manuscript.

## Acknowledgments

We would like to thank Owen White, Anup Mahurkar, and other members of the HMP DACC for helping us make public the revised assemblies of the HMP data. We thank C. Titus Brown and the other reviewers for their thoughtful reviews.

